# Tracking breast cancer progression using Methylscape

**DOI:** 10.1101/2025.07.09.664004

**Authors:** Zhen Zhang, Emtiaz Ahmed, Nicolas Constantin, Jennifer Lu, Darren Korbie, Alain Wuethrich, Abu Ali Ibn Sina, Matt Trau

## Abstract

Cancer progression is intricately driven by epigenetic reprogramming, where aberrant DNA methylation patterns reshape gene regulation and cellular phenotypes. These alterations include promoter hypermethylation of tumour suppressor genes and global hypomethylation, collectively fuelling oncogenesis and disease advancement. We previously introduced the *Methylscape*—a distinct cancer-specific DNA methylation landscape characterized by clustered promoter hypermethylation and gene body hypomethylation—that enhances DNA’s physical affinity for gold surfaces. In this study, we demonstrate that Methylscape can be leveraged to monitor cancer progression. Using a breast cancer epithelial–mesenchymal transition (EMT) model, we observe an increase in Methylscape enrichment of mesenchymal-state DNA during EMT, suggesting that this method can sensitively detect subtle epigenetic remodelling linked to tumour progression. Using a gold-based DNA desorption enrichment strategy coupled with methylation sequencing and qPCR, we show that hypermethylated regions are preferentially enriched. Finally, we developed a low-cost, disposable screen-printed electrode platform for stage-specific breast cancer monitoring. Together, these findings establish Methylscape as a promising biophysical biomarker for non-invasive and real-time monitoring of cancer progression, advancing its potential for clinical translation.

## 1. Introduction

Cancer progression is driven by epigenetic dysregulation, with dynamic changes in DNA methylation emerging as a key mechanism that modulates gene expression and cellular behaviour^1^. Aberrant DNA methylation is characterized by both hypermethylation and hypomethylation, causing disruptions of cellular homeostasis and facilitating oncogenic transformation, and ultimately enabling cancer cells to evade normal regulatory mechanisms and drive uncontrolled growth ^2, 3^. Specifically, hypermethylation tends to cluster at CpG islands within promoter regions, silencing tumour suppressor genes and facilitating oncogenesis. In contrast, global hypomethylation predominantly occurs in intergenic regions, contributing to genomic instability ^2–4^. Together, this unique methylation landscape continues to grow during cancer progression that significantly differs from the normal methylome^5, 6^.

Current methods for monitoring cancer progression include imaging techniques like computed tomography (CT), magnetic resonance imaging (MRI), and positron emission tomography (PET)^7–10^. While these methods provide anatomical insights, they are limited by their inability to detect molecular changes and early-stage progression. Methylation-based methods, including whole-genome bisulphite sequencing (WGBS), reduced representation bisulphite sequencing (RRBS), and targeted methylation assays, are valuable for capturing cancer-specific epigenetic changes ^11, 12^. However, these approaches frequently require intricate workflows, substantial expenses, and extensive sample preparation, limiting their clinical utility. ^13, 14^. Overcoming these challenges is essential for developing simplified, cost-effective, and robust tools capable of reliably monitoring cancer progression at the molecular level, ultimately leading to improved patient outcomes.

Previously, we introduced the concept of *Methylscape*, a phenomenon describing the enrichment of hypermethylated DNA regions through specific methylation changes, including promoter hypermethylation and gene body hypomethylation ^15^. In our earlier work, we demonstrated that DNA derived from a cancerous context exhibited significantly higher physical adsorption on a gold surface compared to DNA from non-cancerous samples ^15^. This was hypothesized to result from the selective exposure of hypermethylated regions, driven by global hypomethylation, which reduces the overall hydrophobicity of the DNA polymer and increases the interaction of methylated regions with hydrophobic surfaces. Exploring this unique surface affinity property of methylated DNA, we developed a simple interfacial biosensing assay capable of electrochemically quantifying DNA adsorption levels and identifying cancer specific methylated DNA.

In this study, we leveraged the unique ability of Methylscape to enrich hypermethylated regions in cancer to demonstrate its potential for tracking epigenetic remodelling during cancer progression (Figure 1). We employed an epithelial-to-mesenchymal transition (EMT) model— a critical process in cancer progression characterised by dynamic DNA methylation changes that promote mesenchymal traits such as motility, invasiveness, and resistance to apoptosis^16,17^. To gain deeper mechanistic insights, a desorption-based approach was developed to extract DNA from the gold surface, followed by qPCR and methylation sequencing. This approach enabled us to determine which specific DNA regions—such as hypermethylated promoters or hypomethylated gene bodies—preferentially interact with the gold surface during EMT progression. This selective isolation of cancer-specific methylated DNA, offers a targeted and efficient approach for analysing epigenetic changes in cancer progression. We also evaluated our method on clinical breast cancer samples and successfully differentiated disease stages using a cost-effective, disposable gold-printed electrode system. The clinical study demonstrates that Methylscape has the potential to serve as a versatile tool for enriching cancer-specific methylation signals, enhancing our understanding of cancer-associated epigenetic changes, and supporting the development of robust, clinically applicable assays for monitoring cancer progression.

**Figure 1.**
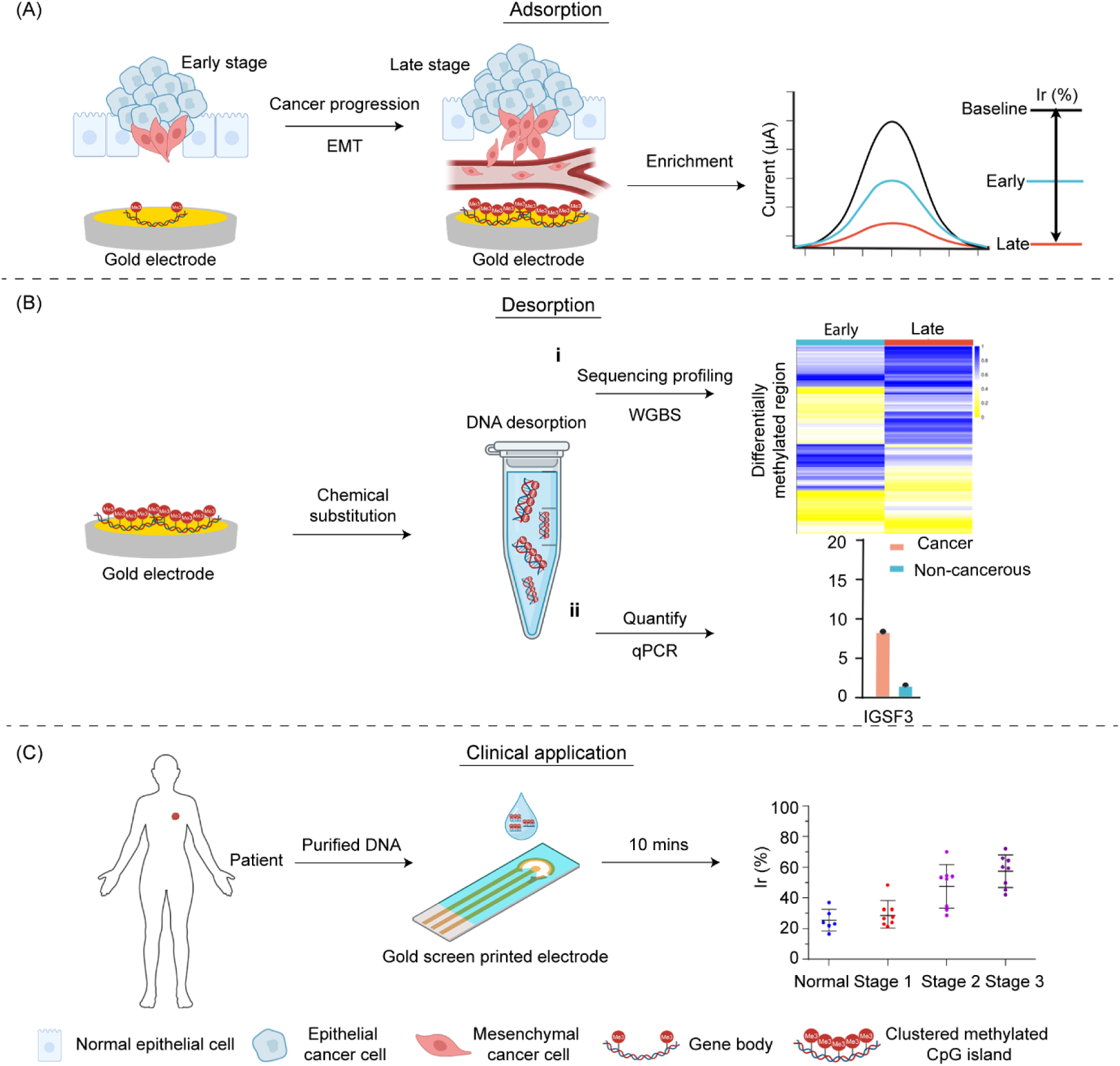
Methylscape analysis for Cancer Progression and Clinical Application. (A) Adsorption: monitoring the EMT process using Methylscape. (B) Desorption: DNA desorption from the gold surface, followed by qPCR and WGBS for comprehensive methylation analysis. (C) Clinical application: employing disposable gold screen-printed electrodes to distinguish between different stages of breast cancer.

## 2. Material and methods

### 2.1 Materials

The epithelial breast cancer cell line MCF-7 (HTB-22) and the mesenchymal cell line MDA-MB-231 (HTB-26) were obtained from ATCC. Cell culture reagents including RPMI1640 medium, fetal bovine serum (FBS), Glutamax, Ultrapure water, penicillin-streptomycin, and Qubit RNA HS Assay Kit were purchased from Thermo Fisher Scientific (USA/Australia), and DNA/RNA concentrations were quantified using the Qubit 4 Fluorometer (Invitrogen, USA). Recombinant TGF-β was from Abcam (UK), and phosphate-buffered saline (PBS; 10 mM, pH 7.4) was purchased from Sigma–Aldrich. Proteinase K (New England BioLabs, p8107s), Buffer AL (QIAGEN, 19075), and the DNeasy Blood and Tissue Kit (QIAGEN, 69504) were used for DNA extraction. For bisulphite conversion, the EZ DNA Methylation-Gold Kit was used (Zymo Research). RNA extraction was conducted using the RNeasy Plus Mini Kit (Qiagen GmbH, Hilden, Germany).

For DNA fragmentation, a Covaris S2 sonicator was used (Covaris), and library preparation for both WGS and WGBS was carried out using the NEBNext Ultra II DNA Library Prep Kit and NEB E7760 or E7595s RNA library kits (New England BioLabs). Sequencing was performed using Illumina platforms: the NextSeq 500 and NovaSeq 6000 systems. Quality control and size distribution analysis were conducted using the Agilent Bioanalyzer (Agilent). Electrochemical measurements were carried out using the CH1040C potentiostat (CH Instruments, USA), and electrodes were fabricated using AZnLOF 2070 photoresist (MicroChem, Newton, MA) and processed with a Temescal FC-2000 e-beam evaporator. Software tools used for primer design and analysis included PrimerROC, PrimerSuite, Trim Galore, and Bismark, along with a custom Python script.

### 2.2 Culture of target cell lines

The selected epithelial breast cancer cell line MCF-7 (ATCCHTB-22) and mesenchymal MDA-MB-231 (ATCCHTB-26) were cultured in RF10 comprising of RPMI1640 culture media (Thermo Fisher Scientific, US) with 10% (v/v) FBS, (Thermo Fisher Scientific, US), 2 mM Glutamax (Thermo Fisher Scientific, US), and 1% penicillin-streptomycin (Thermo Fisher Scientific, US). The cell line was grown in T75 culture flasks in a humidified incubator in 5% CO2 at 37 °C. During Day 0−6, cells were maintained in the low serum culture medium that contained 2% (v/v) FBS. Cells were treated with an additional 10ng·mL−1 TGF-β (Abcam, UK) during the treatment. Cells were passaged every 3 days to maintain consistent confluency (60−70%). The cell line was tested to be mycoplasma-negative before and after treatment. The collected cells from the culture flask were washed and diluted with 1X PBS (10 mm, pH 7.4, Sigma–Aldrich) and DNA was isolated.

### 2.3 DNA extraction and sample preparation

#### 2.3.1 DNA extraction

Collected 10^6^ cells from MCF7 and MDA-MB-231 and washed twice by 1x PBS and stored as pellets in-80 °C. Resuspended the pellet in 200 µL 1x PBS, 20 µL proteinase K (BioLabs®, p8107s) and 200 µL Buffer AL (QIAGEN, 19075), vortexed and incubated at 56 °C for 10 min. Add an equal volume of chloroform: phenol: red: alcohol to the tube and vortexed the solution until the phases are mixed. Centrifuged for 2-3 min at 20238 g to separate phases. Transferred the aqueous phase, added 1/10 of the above volume of 3M Sodium Acetate (pH 5.2), 1 volume of 100% isopropanol. Centrifuged the mixture at 13000g for 15 min at 4 ◦C. Removed the isopropanol and added 1 mL of cold 70% ethanol and mixed the solution by gently inverting it 10 times. Centrifuged the mixture at 13000g for 10 min at 4 ◦C. Removed ethanol and dried the DNA pellet. Added Ultrapure water (Thermo Fisher Scientific, USA) to dissolve the DNA fully. Concentration of DNA measured by Qubit (Qubit 4 Fluorometer, Invitrogen, USA) followed the manufacturer’s instruction.

### 2.4 DNA adsorption on the gold electrode and DPV current measurements

Added readout DNA solution to cover the full electrode gold surface. Measured DPV as baseline under the standard parameters by CH1040C poteniostat (CH Instruments, USA) as the ‘working electrode’. Differential pulse voltametric (DPV) experiments were conducted in 10 mM PBS solution containing 2.5 mM [K_3_Fe(CN)_6_] and 2.5 mM [K_4_Fe(CN)^6^] electrolyte solution. DPV signals were obtained with a potential step of 5 mV, pulse amplitude of 50 mV, pulse width of 50 ms, and pulse period of 100 ms. DPV signals of clean electrodes were measured to get the baseline current and then, added certain concentration of DNA on each electrode and incubated the chip for 10 min at room temperature with 250 rpm vortex. Rinsed the electrodes and dried by nitrogen air. The adsorption competence was measured using the [Fe(CN)_6_]^4-/3-^ redox system. Upon DNA adsorption, the coulombic repulsion between negatively charged ferrocyanide ions in the buffer and negatively charged DNA phosphate groups on the electrode surface partially hinders the diffusion of ferrocyanide ions to the electrode surface. This generates a Faradaic current signal, which is proportionally lower than the bare electrode signals as increasing numbers of DNA molecules become adsorbed onto the surface. The relative adsorption currents (%ir) is measured by the equation:

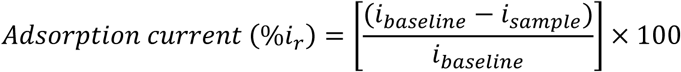

### 2.5 DNA desorption

Following adsorption, the electrodes were washed with 1X PBS buffer three times. To desorb the pre-adsorbed DNA from the gold electrode surface, Mercaptohexanol (MCH) solution was added upon the dried electrode surfaces and incubated for 30 minutes. Since MCH has a stronger affinity for gold than DNA, it effectively displaces the pre-adsorbed DNA molecules from the gold surface upon application. After the incubation period, the solution was collected from the electrode surface and analysed for desorbed DNA using qPCR and electrochemistry.

### 2.6 Detection of desorbed DNA by real-time polymerase chain reaction (qPCR)

Initially, four sets of universal primer-probes of human DNA detection were used for the qPCR assay. The information regarding the universal qPCR assays is summarized in Supplementary Table 1. In the final phase of the study, 8 distinct sets of primers from four different breast cancer genes (such as ALX1, MACF1, IGSF1, and CLPTM1L) were designed to detect the desorbed DNA. Among the primer sets for each gene, 1 primer set (forward and reverse primers) was chosen from the CpG-rich hypermethylated region and another set from the distant gene body region. The sequences of these primers have been included in Supplementary Table 2. qPCR assays were designed using the PrimerROC software and PrimerSuite software packages^18, 19^, together with a custom python script that also predicted the best oligonucleotide to use for fluorescent Taqman assays, targeting an annealing temperature of 60°C.

### 2.7 Sequencing process

#### Whole Genome Sequencing

Whole genome sequencing libraries were constructed by first sonicating the purified desorbed DNA with a Covaris S2 sonicator to achieve a fragment size of approximately 200 bp. The sheared DNA fragments were then subjected to end-repair, A-tailing, and ligation of sequencing adaptors using the NEBNext Ultra II DNA Library Prep Kit (New England BioLabs), following the manufacturer’s recommended protocols. Libraries were then quantified using a Qubit fluorometer (Thermo Fisher Scientific) and diluted to the required concentration. Sequencing was conducted on an Illumina NextSeq 500 platform using a 150 MId kit, with a paired end 2×71 bp read configuration and dual index adaptors to enable accurate demultiplexing of samples. Quality control of sequencing reads was performed post-run to assess read quality.

#### Whole Genome Bisulphite sequencing

Whole-genome bisulphite sequencing libraries were prepared to assess genome-wide DNA methylation patterns. Purified desorbed DNA was fragmented to an average size of ∼200 bp using a Covaris S2 sonicator, followed by end-repair, A-tailing, and adaptor ligation using the NEBNext Ultra II DNA Library Prep Kit for Illumina (New England BioLabs), according to the manufacturer’s protocols. Adaptor-ligated DNA was treated with the EZ DNA Methylation-Gold Kit (Zymo Research) to convert unmethylated cytosines to uracils, while methylated cytosines remained unchanged. The bisulphite-treated DNA was then amplified with high-fidelity, low-bias PCR to create the final library. Quality and size distribution of the libraries were verified using an Agilent Bioanalyzer, and libraries were quantified via qPCR. Sequencing was carried out on an Illumina NovaSeq 6000 system in paired-end 150 bp mode. Raw sequence reads were processed using Trim Galore to remove adaptor sequences and low-quality bases, followed by alignment to a bisulphite-converted reference genome using Bismark. Bisulphite conversion efficiency was assessed using spike-in unmethylated lambda DNA.

### 2.8 RNA extraction

High-quality genomic RNA was extracted from cultured breast cancer cells using Qiagen RNeasy Plus Mini Kit (Qiagen GmbH, Hilden, Germany) in accordance with the manufacturer’s protocols. Moreover, we used the Qubit 4.0 Fluorometer (ThermoFisher, Australia) to measure the concentration and quality of the extracted RNA using the Qubit RNA HS Assay Kit (ThermoFisher, Australia). Depending on sample volume, the assay kit provides an accurate measurement for initial RNA sample concentrations of 0.2 to 200 ng/µL, providing a detection range of 4−200 ng.

### 2.9 RNA sequencing

The RNA library was prepared with NEB E7760 kit for rRNA depleted FFPE RNA and NEBNext Ultra II (E7595s) Directional RNA library prep kit for Illumina. RNA Sequencing was performed on the NextSeq 500 Sequencer using default parameters.

### 2.10 RNA-seq and enrichment analysis

The Tuxedo suite (adapted from Pertea et al (Nature Protocol, 2016)) was used to perform transcript-level expression analysis of RNA-seq experiments with hist2, stringtie, and ballgown. Briefly, hisat2 (2.2.1) was used to map fastq files that were previously trimmed with [trimmomatic?] were mapped to the hg38 genome (options: hisat2-p 6--dta--sensitive--no-soft-x <hisat2.index.prefix>-q-1 sample.R1.fastq-2 sample.R2.fastq-S sample.sam). Transcripts were assembled based on mapping to genome using the stringtie (2.1.6) package (option:--merge-p 6-G reference.gtf-o merged_gtf_file merged_file). gffcompare (0.112) was used to compared the assembled transcripts to known transcripts. stringtie (2.1.6) was used to estimate abundance of assembled transcripts (options:-e-B-p 6-G merged_gft_file-o ballgown_gtf_file sample.sortedbam). Transcript annotation and gene level expression analysis was conducted using the R package (ballgown) and visualized using ggplot2. Differential expression analysis was performed using the R package EnhancedVolcano using standard protocols outlined in Blighe (2018), where p ≤ 0.05, and fold change was set to 1. Enrichment analysis was performed in R where transcripts that displayed significant fold change (FDR <= 0.01, and log2 fold change >= 1) were selected for enrichmenta. (All scripts for differential analysis and enrichment analysis are available on request).

## 3. Results and Discussion

### 3.1 Application of Methylscape for Monitoring Cancer Progression via the EMT Model

Epithelial-mesenchymal transition (EMT) is a critical process in cancer progression that plays a key role in initiating metastasis. During EMT, epithelial cancer cells undergo significant phenotypic and morphological changes, transforming into mesenchymal-like cells. This transition enables the cells to acquire enhanced migratory and invasive properties, leading to the release of circulating tumour cells (CTCs) into the bloodstream, which can subsequently result in cancer metastasis^16, 20, 21^. During metastasis, the EMT process enables CTCs to evade immune surveillance, contributing to the development of drug resistance. Interestingly, DNA methylation-mediated dynamic epigenetic reprogramming is a key mechanism that initiates the EMT process^17, 22–25^. For instance, during the EMT process, dynamic changes in methylation patterns in the promoter regions of epithelial genes (e.g., CDH1) and mesenchymal genes (e.g., VIM) in response to EMT stimuli such as TGF-β, hypoxia, growth factors from the microenvironment^26^. Moreover, recent reports have found that DNA methylation changes underlying EMT are among the most common mechanism leading to the development of therapy resistance^17^. Since Methylscape is designed to identify changes in the DNA methylation landscape in cancer, we decided to use a synthetic EMT model to determine whether our method can accurately detect cancer EMT events and trace cancer progression in a breast cancer model.

To test our hypothesis, we treated two breast cancer cell lines with TGF-β to create a synthetic EMT-induced model. The epithelial breast cancer cell line (MCF7) and the mesenchymal breast cancer cell line (MDA-MB-231) were cultured for 12 days: 6 days with TGF-β treatment and 6 days post-treatment. Cells were collected at six time points: Day 0 (pre-treatment), Day 3 and Day 6 (during treatment), and Day 9 and Day 12 (post-treatment). The EMT transition was confirmed by flow cytometry (**Figure 2A** and **ESI Figure S1A**), showing a downregulation of E-cadherin and an upregulation of N-cadherin on Day 3 and Day 6, indicating a shift to a mesenchymal phenotype. After the treatment was withdrawn, both marker expressions returned to a more epithelial-like phenotype, suggesting that the EMT process was reversed after the cessation of treatment.

**Figure 2.**
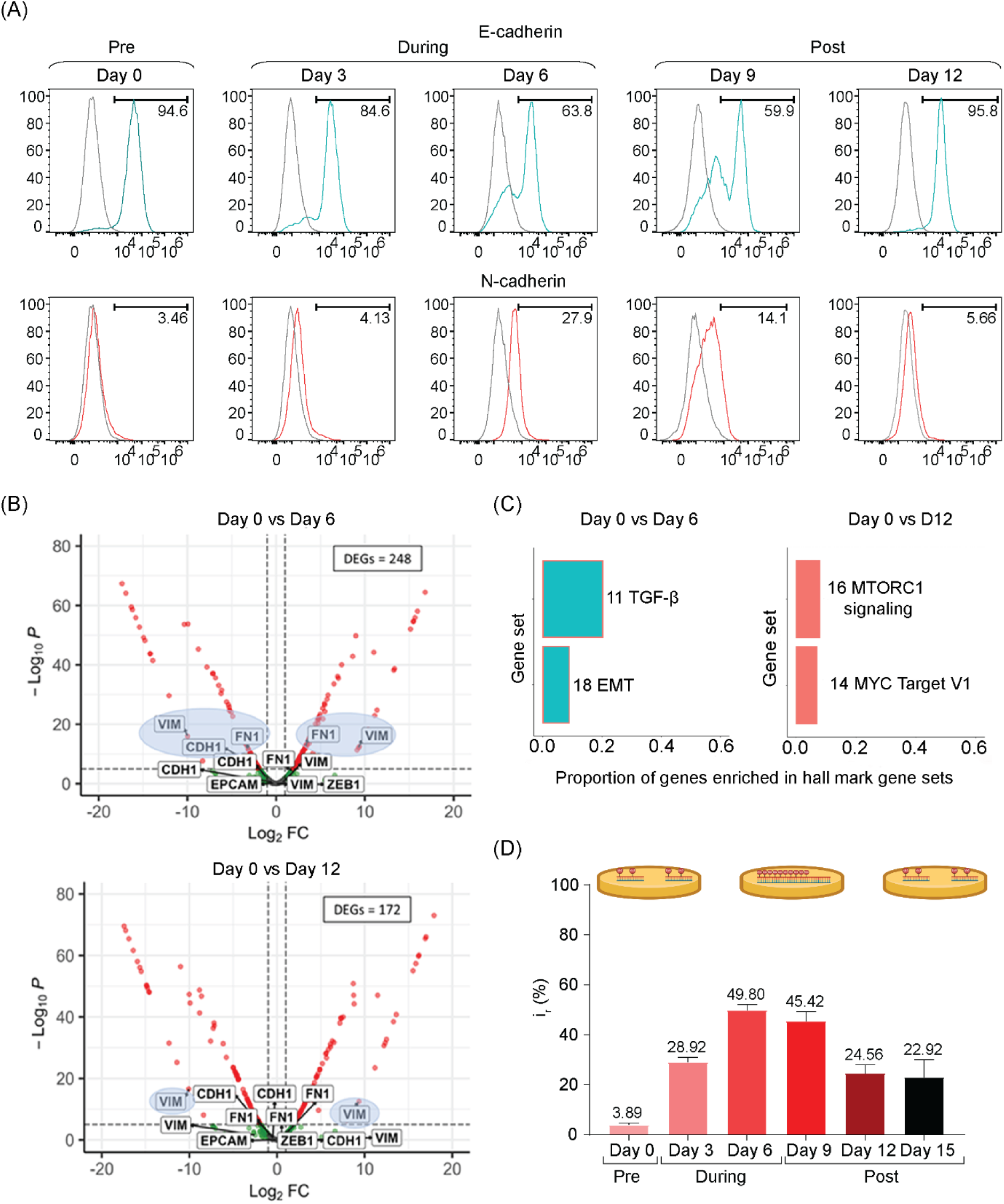
Methylscape analysis on drug induced EMT model (MCF7) for investigating cancer progression. (A) Flow cytometry analysis of E-cadherin and N-cadherin in epithelial breast cancer cell line MCF7 among TGF-β treatment pre-treatment (Day 0), during treatment (Day 3 & Day 6), post-treatment (Day 9 & Day 12). (B) The number of differentially expressed genes (DEGs) (FDR <0.05 and |log2fold change| ≥ 1) in the epithelial cell model (MCF7) during-and post-treatment were compared to the pre-treated samples. (C) Enrichment analysis of differentially expressed transcripts during-and post TGF-β treatment on the epithelial cell line (MCF7). The number of enriched transcripts is shown next to each plot while the y axis displays the proportion of differentially expressed transcripts that are enriched in each pathway when compared with the hallmark gene sets. The number of enriched transcripts is shown next to each plot while the y axis displays the proportion of differentially expressed transcripts that are enriched in each pathway when compared with the hallmark gene sets. (D) Methylscape analysis for adsorbing MCF7 DNA on gold surface among the timepoints.

To evaluate EMT-associated transcript changes, we conducted RNA sequencing analysis on MCF7 cells at pre-treatment (Day 0), during treatment (Day 6), and post-treatment (Day 12),using EdgeR to identify differentially expressed genes (DEGs). The DEGs were visualized in a volcano plot, with significance thresholds set at a p-value ≤ 0.05 and a log10 fold change of 1. While several EMT-related genes, including CDH1, EPCAM, VIM, and ZEB1, were expressed during TGF-β treatment, only FN1 and VIM were significantly differentially expressed, corresponding to the mesenchymal markers fibronectin and vimentin, respectively (**Figure 2B**). After treatment withdrawal, FN1 expression decreased, and CDH1, which was downregulated during treatment, returned to baseline levels, indicating a reversal of the EMT process (**Figure 2C**). To further investigate, we analysed the mesenchymal cell line MDAMB231 under similar conditions, where TGF-β was used to enhance the EMT transition. Consistent with MCF7, VIM and FN1 were differentially expressed during and after treatment, while CDH1 was downregulated six days post-treatment (**ESI Figure S1B & C**). This mirrored the pattern observed in MCF7 cells, supporting the induction of EMT by TGF-β treatment.

To understand the potential functions of the differentially expressed EMT-associated transcripts, we performed a hypergeometric enrichment analysis on genes with significant fold changes during and after TGF-β treatment. In MCF7 cells, the analysis revealed enrichment in the TGF-β signalling pathway during treatment, consistent with the induction of a mesenchymal state (**Figure 2C**). However, this enrichment disappeared after treatment was withdrawn, aligning with the observed phenotypic reversal (**Figure 2C**). Similarly, in MDAMB231 cells, differentially expressed transcripts were enriched in the TGF-β signalling and EMT pathways during treatment, with this enrichment dissipating post-treatment (**ESI Figure S1C**). These findings highlight the role of TGF-β in driving EMT, as evidenced by the transient activation of these pathways during treatment.

Lastly, after validation of the drug-induced EMT model, we aim to utilize Methylscape for monitoring the EMT process. We optimized the DNA adsorption condition on a gold surface by testing different DNA concentrations and solvation conditions. Four DNA concentrations (1 ng/µL, 2 ng/µL, 5 ng/µL and 10 ng/µL), and two solvation conditions (5X SSC buffer and H_2_O) were evaluated. As shown in **ESI Figure S2**, we observed increased DNA adsorption on the gold surface with rising DNA concentrations at three treatment points. DNA concentrations of 5 ng/µL and 10 ng/µL resulted in saturated measurements on the gold surface, particularly during mid-treatment (Day 6) and post-treatment (Day 12). In contrast, the 1 ng/µL DNA samples produced weaker signals compared to the 2 ng/µL samples. Based on these findings, we selected 2 ng/µL as the standard DNA concentration.

We also observed higher DNA adsorption on the gold surface when DNA was dissolved in 5X SSC buffer compared to pure H_2_O. According to a previous study^15^, the addition of salt from the 5X SSC buffer can induce charge neutralisation, thereby promoting DNA adsorption on the gold surface—results consistent with our observations. However, monitoring the EMT process requires a more sensitive detection method, and we needed to minimize signal saturation caused by salt-induced charge neutralisation. When we switched the solvent to H_2_O, we observed lower signals compared to 5X SSC. For example, in **ESI Figure S2**, MCF7 DNA in 5X SSC buffer maintained high adsorption at Day 12 post-TGF-β treatment, but when the same DNA sample was dissolved in pure H_2_O, DNA adsorption decreased on Day 12 (**ESI Figure S2**). This result aligns with previous flow cytometry findings, which showed a restoration of the epithelial phenotype on Day 12, consistent with the decreased adsorption observed in H_2_O on Day 12. This phenomenon suggests that DNA adsorption induced by the hypertonic solution could lead to false-positive measurements during the EMT process. Therefore, the 2 ng/µL DNA concentration with H_2_O was selected as the standard condition for subsequent experiments.

Based on the optimization assay described earlier, we proceeded with 2 ng/µL of DNA dissolved in pure H_2_O as our standard condition. Both breast cancer cell lines showed a significant increase in DNA adsorption during the TGF-β treatment period (Day 3 and Day 6) (**Figure 2D and ESI Figure S1D**). After the treatment was withdrawn, a decrease in DNA adsorption was observed in both cell lines, which was consistent with the flow cytometry results. These findings demonstrate that Methylscape can monitor the EMT process by tracking DNA adsorption on the gold surface, with results that are further corroborated by transcriptomic changes detected by RNA sequencing and phenotypic changes observed through flow cytometry. These results suggest that Methylscape could serve as an advanced tool for monitoring cancer progression by tracking changes in its levels.

While these results are promising, the precise mechanism behind Methylscape’s ability to detect EMT events, particularly its interaction with hypermethylated promoters and hypomethylated gene bodies on a gold surface, remains to be understood. To address this, we thought to develop a method to desorb untreated and treated DNA from the gold surface, followed by qPCR and WGS, to gain a deeper understanding of the Methylscape mechanism.

### 3.2 Understanding the Mechanism of Methylscape Through the Desorption Model

Clustered hypermethylation at DNA promoter regions is a key factor in cancer progression, as it silences transcription and inactivates crucial genes. This epigenetic reprogramming alters the physicochemical properties of the cancerous genome, significantly impacting its interaction with gold surfaces. To gain deeper mechanistic insights, our study developed a desorption model to investigate the fundamental interactions between DNA and gold surfaces, demonstrating that hypermethylated cancerous DNA exhibits a stronger adsorption preference compared to non-cancerous DNA. **ESI Figure S3** represents the scheme of the DNA adsorption, desorption, qPCR detection, and sequencing approaches. Briefly, purified DNA from the MCF7 breast cancer cell line was initially adsorbed onto gold electrode surfaces and relative adsorption was measured electrochemically (**ESI Figure S4 A and C**). Desorption of DNA was then achieved either electrochemically or via a competitive method using MCH, which has a higher affinity for gold than DNA and effectively displaced the adsorbed DNA. The MCH-based competitive desorption method was selected for subsequent experiments due to its superior yield. The viability of the desorbed DNA for downstream analysis was confirmed by PCR amplification **(ESI Figure S4 B and D**) leveraging a universal primer-probe sets (n = 4) to detect any human DNA **(Supplementary Table 1).** Finally, the methylation analysis of the desorbed DNA was performed by qPCR, whole genome sequencing and whole genome bisulphite sequencing approaches.

To obtain greater insight into the desorbed DNA, we designed 8 distinct sets of primers from four different genes (such as ALX1, MACF1, IGSF3, and CLPTM1L) to detect the desorbed DNA more specifically. For each of the genes, 1 primer set (forward and reverse primers) was chosen from the CpG-rich hypermethylated promoter region and another set from the distant gene body (intergenic) regions which are hypomethylated **(Supplementary Table 2).** These genes were selected based on TCGA (The Cancer Genome Atlas) data, which revealed that they exhibit promoter hypermethylation and gene body hypomethylation in breast cancer. **ESI Figure S5** presents the qPCR results, showing successful amplification of all targeted regions, thereby confirming the adsorption of hypermethylated and gene-body regions of DNA onto the gold electrodes.

To investigate further, we analysed the desorbed DNA by qPCR to calculate copy numbers and concentration of the desorbed DNA sample to reveal the methylated DNA enrichment ratio (i.e., DNA copies from promoter regions versus intergenic regions). **Figures 3A and ESI Figure S6** show the computed copy numbers and concentrations of desorbed MCF-7 DNA detected by primers targeting the promoter and gene body regions of each gene, respectively. Both copy numbers and concentrations of DNA were higher in promoter regions where the IGSF3 promoter region detected 13 copies/μL, while the gene body region detected 4 copies/μL, yielding a promoter/gene-body enrichment ratio of 8.33. Similar ratios were observed for ALX1, MACF1, and CLPTM1L, at 1.40, 3.06, and 4.39, respectively (**Figure 3B**). These findings support the hypothesis that hypermethylated regions preferentially adsorb onto the gold surfaces. Furthermore, Figures 3A and 3B illustrate the copy number and promoter-to-gene body enrichment ratio of desorbed DNA from cancerous (MCF7) and non-cancerous (healthy buffy coat) samples. As anticipated, the healthy samples exhibited lower copy numbers and promoter-to-gene body ratios compared to the MCF7 DNA, reflecting reduced desorption yields due to weaker adsorption onto the gold surface. These findings are consistent with our previous observations that normal genomic DNA—despite its higher global methylation levels (60–80%)—shows reduced adsorption to gold. This is likely due to the hydrophobicity induced by widespread methylation, which limits the accessibility of locally clustered hypermethylated regions to interact effectively with the gold surface.

**Figure 3.**
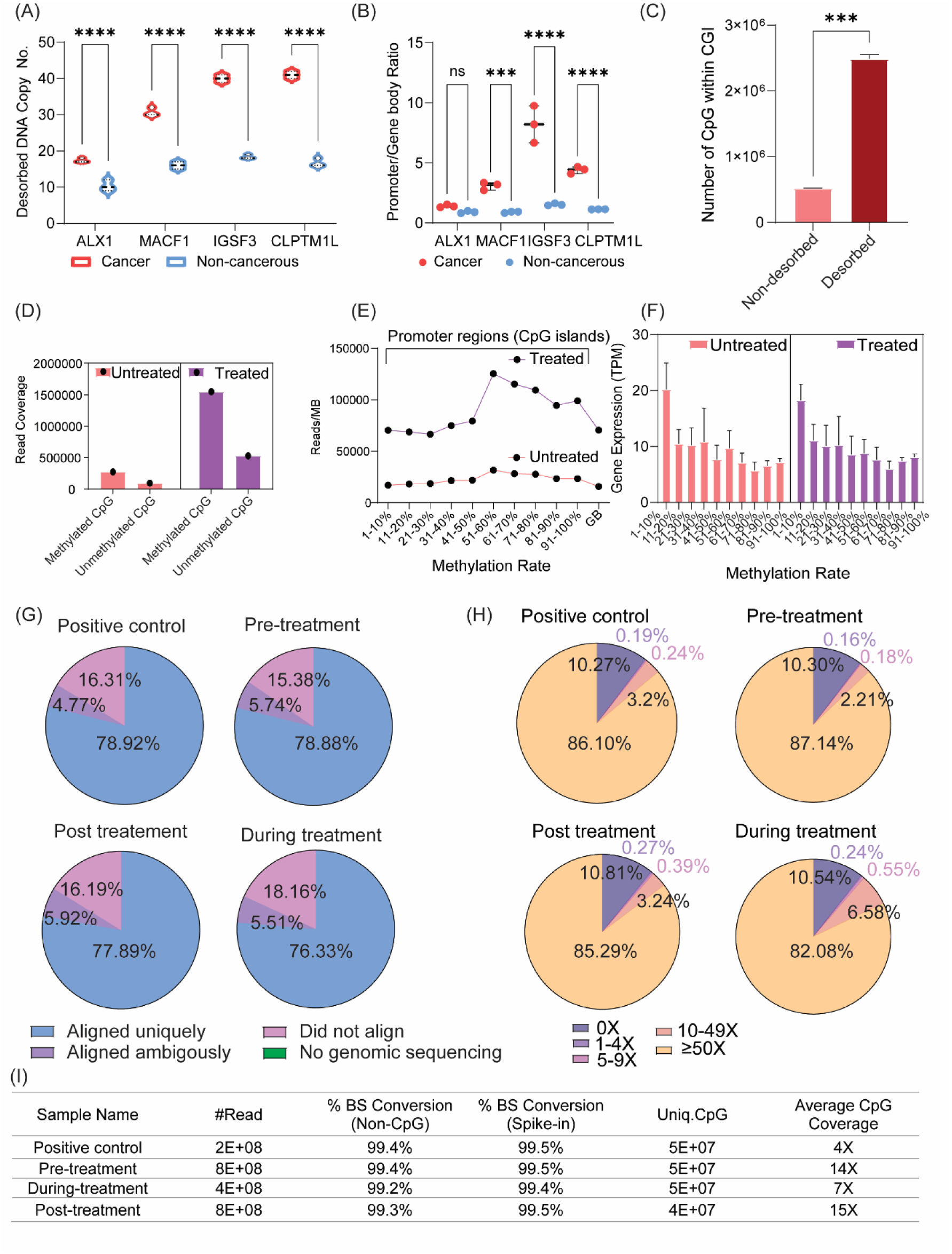
qPCR and Whole-Genome and Whole Genome Bisulphite Sequencing of Desorbed DNA. (A) qPCR analysis revealed a significantly higher DNA copy number and (B) methylated DNA enrichment ratio in MCF-7 desorbed DNA compared to non-cancerous DNA. (C) A highly significant difference was observed in the number of CpGs within CpG islands (CGIs) between non-desorbed and desorbed DNA. (D) Methylated DNA coverage at promoter CpG islands was substantially higher than unmethylated DNA coverage in both untreated and TGF-β-treated samples. (E) An increasing trend in read coverage per MB correlated with a higher methylation ratio (%) in both untreated and treated desorbed DNA samples at promoter regions. Additionally, methylated read coverage at promoter regions was significantly higher than at intergenic regions for both sample types. (F) Average gene expression showed a decreasing trend with increasing methylation ratio (%) at promoter regions in both untreated and treated RNA samples. (G) Most reads (∼76–79%) aligned uniquely, ensuring high sequencing quality. (H) Majority of CpG islands had high sequencing depth (≥50X), supporting robust methylation analysis. (I) High bisulphite conversion efficiency (∼99.2–99.5%), stable CpG coverage (∼45–48 million CpGs), and adequate sequencing depth (4X–15X) confirm data reliability.

This findings from the qPCR assays prompted us to sequence the desorbed DNA to further investigate the mechanism of Methylscape. We performed whole genome sequencing analysis of two desorbed DNA samples (untreated and TGF-β treated Day 6 MCF7 DNA) and compared read coverage between promoter regions (CpG islands) and intergenic regions (**ESI Figure S7**). In our whole genome sequencing analysis, we identified a significant difference in the number of CpG sites within CpG islands (CGIs) between non-desorbed (positive control) and desorbed untreated MCF7 DNA samples. The desorbed DNA displayed a substantially higher number of CpG sites in CGIs compared to the non-desorbed sample (**Figure 3C**). This is potentially due to the preferential adsorption of hypermethylated CpG-rich DNA on the gold surface. These findings indicate that desorbed DNA provides a selective enrichment of methylated DNA within CpG islands, which could be beneficial for accurately characterising epigenetic modifications in cancer cells. To determine the methylation status of the sequenced reads, we used TCGA methylation data from breast cancer patients and correlated it with our sequenced reads, assuming similar methylation pattern in MCF-7 breast cancer cells. **Figure 3D** shows that read coverage of methylated CpGs was significantly higher than that of unmethylated CpGs in promoter regions for both samples, supporting the hypothesis that methylated DNA has a higher affinity for gold surfaces, consistent with a recent simulation study ^27^. Interestingly, the read coverage of both methylated and unmethylated CpGs are significantly higher in the case of TGF-β treated Day 6 MCF7 Samples (Mesenchymal states) which is consistent with the higher adsorption of mesenchymal DNA towards the gold surface. This desorption sequencing data supports the higher adsorption of DNA from TGF-β-treated Day 6 MCF7 cells onto the gold surface (Fig. 2D). This further confirms that Methylscape can effectively track the dynamic changes in methylation patterns during EMT and cancer progression— the gradual increase in CpG hypermethylation and intergenic hypomethylation.

To better understand the methylation status of desorbed DNA samples, we created a graph categorizing promoter regions by percentage of methylation (0-10%, 11-20%, 21-30%, etc.) and assessed whether read coverage increased with higher methylation percentages. We also compared methylation rates between promoter and gene body (GB) regions. **Figure 3E** shows that read coverage increased with higher methylation percentages in promoter regions for both samples, particularly for the TGF-β-treated sample. Additionally, methylated DNA read coverage was significantly higher in promoter regions (CpG islands) compared to intergenic regions (gene bodies). Read coverage was normalized per megabase (MB) due to significant differences in nucleotide base counts among the methylation sub-groups. Moreover, we sequenced RNA from untreated and TGF-β treated MCF-7 cells to explore the link between methylation status and gene expression. **Figure 3F s**hows that average gene expression decreased as methylation levels increased in promoter regions. This supports previous findings that hypermethylation can inhibit gene expression and regulate transcription. Methylation recruits’ methyl-CpG-binding domain proteins and histone deacetylases, which prevent RNA polymerase binding and repress gene expression^4, 28^.

To directly investigate the epigenetic reprogramming patterns, we performed whole-genome bisulphite sequencing (WGBS) on a set of three desorbed MCF7 DNA samples (untreated/pre-treated Day-0 MCF7, TGF-β-treated Day-6 MCF7, and post-treated Day-12 MCF7). The Bismark alignment score from WGBS were present in **Figure 3G**, revealing that approximately 78% of reads are uniquely aligned. This high alignment rate indicates excellent alignment quality and supports the reliability of our sequencing data for accurate methylation analysis. Moreover, our WGBS experiments achieved high CpG island coverage in all samples (positive control, pre-treatment, during treatment, and post-treatment), with over 50x coverage for around 25,000 CpG islands (**Figure 3H)**. This coverage ensures a robust and confident assessment of methylation patterns within CpG-rich regulatory regions, facilitating precise comparative analyses of methylation changes across different treatment stages. **Table 3I** evaluates sequencing quality, demonstrating high bisulphite conversion efficiency (99.2– 99.5%), a critical measure of reliable unmethylated cytosine deamination.

Next, we performed methylation Analysis of our WGBS data and identified differentially methylated regions (DMRs) and differentially methylated cytosines between the three desorbed DNA conditions: untreated, treated, and post-treated samples. **Figure 4 A(i), B(i) and Table 4C** shows that there are higher number of hypermethylated regions and methylated cytosines in the desorbed TGF-β-treated Day-6 MCF7 DNA samples in comparison to the desorbed untreated Day 0 DNA samples. The WGBS data from the desorbed DNA further validates our earlier qPCR and whole-genome sequencing findings, reinforcing that hypermethylated regions exhibit a stronger affinity for the gold surface and as cancer progresses, the adsorption of hypermethylated DNA increases—a trend that Methylscape can effectively identify. Figures 4A(ii–iii) and 4B(ii–iii) present the DMRs and DMCs observed under additional experimental conditions.

**Figure 4.**
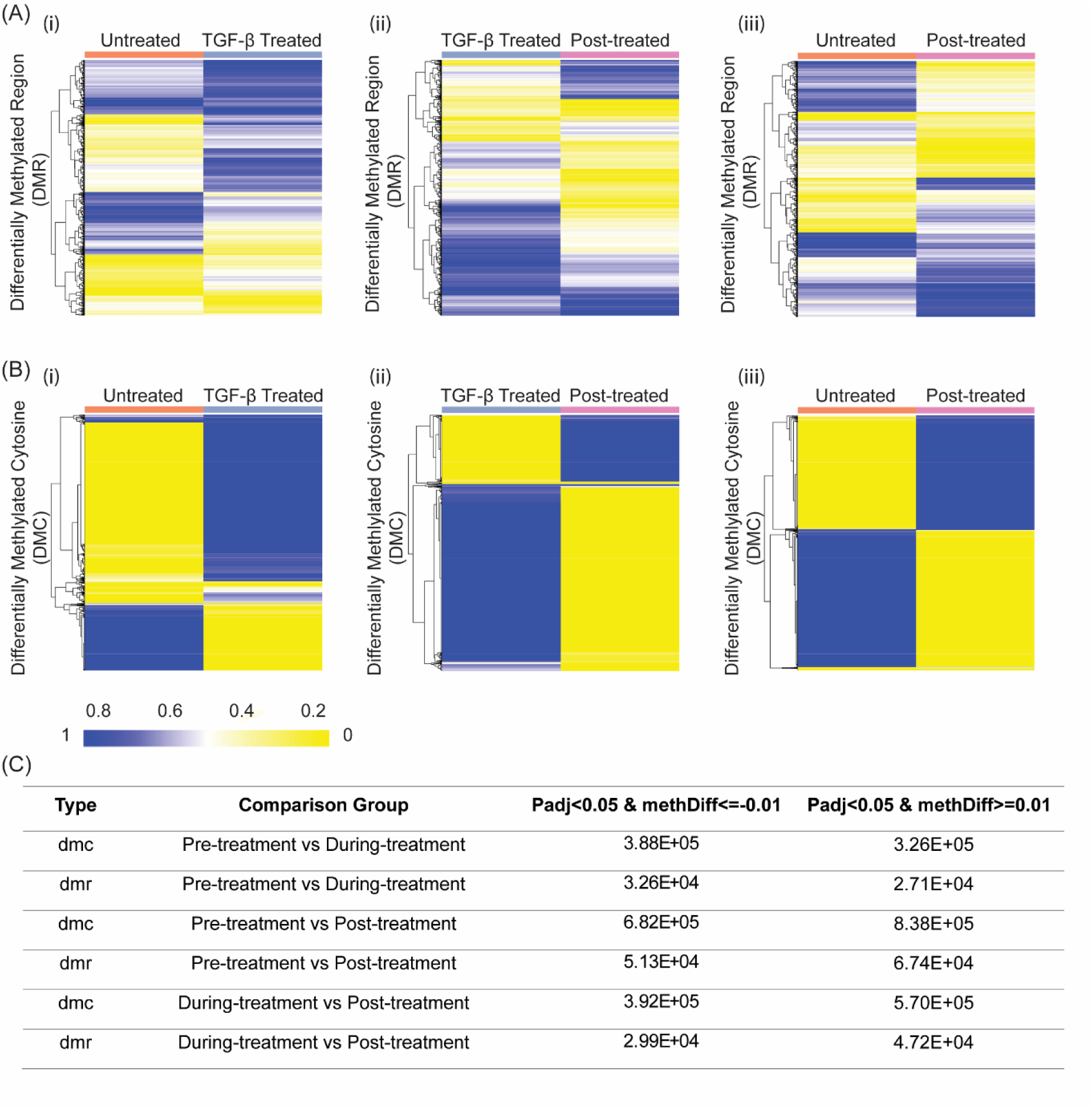
The WGBS data reveal treatment-induced DNA methylation changes across three conditions: untreated, treated, and post-treated samples. Differentially methylated regions found in untreated vs treated samples (Ai), Treated vs Post-treated (Aii) and Untreated vs Post-treated samples (Aiii) and differentially methylated cytosines found in untreated vs treated samples (Bi), Treated vs Post-treated (Bii) and Untreated vs Post-treated samples (Biii) and Differentially methylated cytosines. (C) Table showing the actual number of DMCs and DMRs were detected across treatment conditions.

### 3.3. Clinical translation - Methylscape diagnostic assay using disposable electrodes to detect breast cancer progression

After validating Methylscape for monitoring the EMT process and investigating its underlying mechanisms using a desorption enrichment method combined with sequencing, we aimed to assess the translational potential of the Methylscape platform in tracking cancer progression. Leveraging the unique physicochemical properties of cancer DNA, which preferentially adsorbs to a bare gold surface, we evaluated epigenome remodelling in a cohort of breast cancer patients across different stages (I–III).

Considering that clinical pathology prefers single-use detection test to minimise sample contamination and operator variability, one potential avenue for electrochemical biosensing-based diagnostic application is screen-printed electrodes (SPE) as they offer the advantages of low-cost production and practical feasibility. The presented interfacial biosensing methodology was thus developed on commercially available gold SPE by comparing the relative DNA adsorption to each electrode surface during resistive measurement using DNA derived from a breast cancer model cell line (MCF7) to simulate tumour sample and buffy coat DNA from a pool of blood donors to simulate a control, healthy sample. All genomic DNA samples used in this exploratory study were purified using a gold-standard phenol-chloroform extraction process and for each electrode testing, 100 ng (i.e., 10 μL at 10 ng/μL) of DNA was incubated on the gold surface for 10 minutes before differential pulse voltammetry (DPV) experiment.

Results in **Figure 5A** shows the ability of the developed SPE biosensing approach to differentiate between control buffy coat and MCF cancerous samples with a difference in gold adsorption of almost 50%, a nearly four-fold increase compared to conventional re-usable lab-based electrode system.

**Figure 5:**
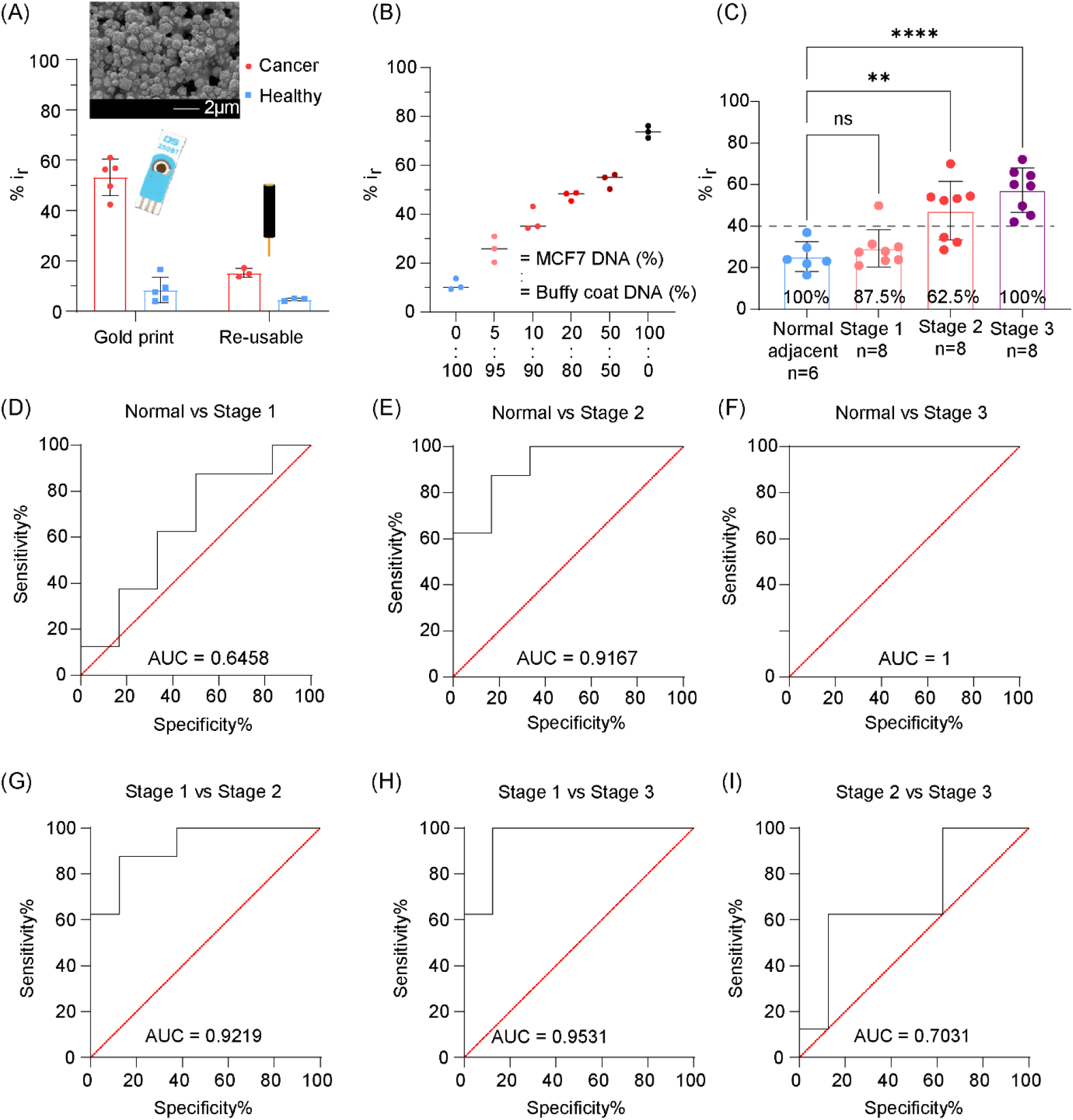
Clinical utility of the screen-printed gold electrode (SPE) platform to stratify stages of breast tumours using Methylscape. (A) Differences in biosensing detection performance between single-use (SPE) and traditional re-usable gold electrodes. (B) Graphs showing the relative gold adsorption (%Ir) of pooled buffy coat DNA samples spiked with MCF7 cancer DNA at different ratios (from 0% to 100%). (C) Clinical analysis of DNA purified from 6 different normal adjacent breast tissues (blue) and 24 different breast tumour tissue clinically classified as stage 1 (pink, n = 8), stage 2 (red, n = 8) and stage 3 (purple, n = 8). Statistical significance between groups (unpaired t-test) is shown for normal adjacent vs. stage 1 (p = 0.387, ns), vs. stage 2 (p = 0.046, **) and vs. stage 3 (p < 0001, ****) respectively.

Given that many tumour samples contain significant proportions of normal adjacent tissue, the sensitivity of the proposed electrochemical detection system to contaminating “normal” DNA was evaluated by spiking buffy coat DNA with MCF7 breast cancer line DNA at different ratios. Results in **Figure 5B** indicate that the SPE platform can detect as low as 5% of cancer DNA in a matrix of 95% of buffy coat DNA when analysing 100 ng of samples.

Finally, to demonstrate the clinical utility of the methodology to detect global epigenetic remodelling associated with cancer progression, gDNA purified from breast cancer tumour samples across three different cancer stages (i.e., stages 1, 2 and 3) was profiled and compared to gDNA isolated from normal adjacent breast tissues. Eight different patients were selected for each stage for a total of 24 breast cancer samples, together with six normal adjacent breast tissue controls (see **Supplementary Table S3** for patient details). Each tumour DNA samples were analysed in electrode triplicates while normal adjacent samples in duplicates. The resulting data show a global trend with the relative DNA gold adsorption increasing with tumour stages, where stage 2 and 3 samples tend to sit above an arbitrary chosen threshold value of %Ir = 40% as opposed to stage 1 and normal adjacent samples (**Figure 5C**). More precisely, when considering the mean %Ir value of each patient sample and comparing each different diseased group, 100% of stage 3 samples (8/8) and 62.5% of stage 2 samples (5/8) were above the 40% threshold, while 87.5% of stage 1 samples (7/8) and 100% of normal adjacent samples (6/6) were below. In addition, statistical significances (unpaired t-test) were observed when comparing the patient’s data from the normal adjacent group to the ones from stage 2 (p-value = 0.0046) and stage 3 (p-value < 0.0001) respectively. However, no significant statistical difference was observed between the normal adjacent and the stage 1 samples (p = 0.387). Together, these data translate to an area under the receiving operating curve (AUROC) of 0.65, 0.92, and 1 when comparing normal samples vs stage I, normal vs stage 2, and normal vs stage 3 samples respectively (Figure 5D-F), and to an AUROC of 0.92, 0.95, and 0.70 when comparing stage 1 vs2, stage 1vs 3, and stage 2 vs 3 (**Figure 5G-I**).

These promising results reveal the potential of the presented Methylscape to stratify stages of the disease by tracking evolving epigenomes associated with cancer progression. As opposed to other existing approaches in the field, our unique methodology focuses on the analysis of the physical consequences induced by epigenetic rearrangements of DNA molecules. Such features are difficult to detect by sequencing analysis only and could provide complementary information to conventional cancer diagnostic technologies for a more comprehensive diagnosis. Given the clinical potential and the unprecedented simplicity of the presented SPE platform (e.g., label-free and functionalisation-free sensors, no DNA treatment nor amplification), it is believed to find great opportunities as a cost-effective and rapid DNA methylation-based cancer detection assay to screen population in low resource settings.

The horizontal dotted line shows an arbitrary threshold value of %Ir = 40%. Per cent values (%) indicate the proportion of patient samples with a mean %Ir value being below the %Ir = 40% threshold for normal adjacent and stage 1 or being above the threshold for stages 2 and 3. (D-I) Receiver operative characteristics curves were computed using the data plotted in (C).

## 4. Conclusion

Our study demonstrates that Methylscape offers a simple, rapid, and cost-effective strategy for monitoring cancer progression by quantifying DNA adsorption onto a gold surface. Using an EMT-based cell line model, we first validated the concept, showing significantly higher Methylscape signals in DNA from EMT-transitioned cells. To gain mechanistic insights, we desorbed the pre-adsorbed DNA from the gold surface and performed qPCR, whole-genome sequencing, and whole-genome bisulphite sequencing. The results revealed preferential adsorption of clustered hypermethylated DNA regions onto gold, supporting our hypothesis that hypermethylated domains exhibit stronger gold affinity than hypo-or unmethylated regions. To facilitate clinical translation, we employed a disposable screen-printed electrode as a low-cost detection platform. Together, these findings highlight Methylscape’s potential as a robust tool for tracking cancer progression, enriching cancer-specific methylated DNA, and enabling the development of clinically applicable assays for disease monitoring.

## Abbreviations

DNA: (Deoxyribonucleic acid)
RNA: (Ribonucleic acid)
Dnmts: (DNA methyltransferases)
SAM: (S-adenyl methionine)
DPV: (Differential Pulse Voltammetry)
qPCR: (Real-time Polymerase Chain Reaction)
WGS: (Whole Genome Sequencing)
TGF-β: (Transforming Growth Factor-Beta)
MCF-7: (Michigan Cancer Foundation-7)

## CRediT authorship contribution statement

**Zhen Zhang:** Investigation, Methodology, Writing – original draft. **Emtiaz Ahmed:** Investigation, Methodology, Writing – original draft. **Nicolas Constantin:** Investigation, Methodology, Writing – original draft. **Jennifer Lu:** Methodology, Writing – review & editing. **Alain Wuethrich:** Methodology, Writing – review & editing. **Darren Korbie**: Methodology, Supervision, Writing – review & editing**. Abu Ali Ibn Sina:** Conceptualization, Supervision, Funding acquisition, Writing – review & editing. **Matt Trau:** Conceptualization, Supervision, Funding acquisition, Writing – review & editing.

## Data availability

Data will be made available on request.

## Acknowledgments

This work was supported by NHMRC Investigator Grant (APP1175047 for AAIS; APP2034488 for AW), Australian Research Council Discovery Project (DP180102836 for MT), and Cancer Australia (2010799 for MT and AW). These grants significantly contributed to the environment to stimulate the research described here. EA thanks the support from Australian Government Research Training Program (RTP) Scholarship (The University of Queensland) to EA. The gold electrodes fabrication works were conducted at the Queensland node of the Australian National Fabrication Facility (Q-ANFF).

## Appendix A. Electronic Supporting Information (ESI)

The supporting data to this article has been attached.

